# Nitric oxide (NO) signaling in *Trichoplax* and related species: Microchemical characterization and the lineage-specific diversification

**DOI:** 10.1101/2020.04.10.034207

**Authors:** Leonid L. Moroz, Daria Y. Romanova, Mikhail A. Nikitin, Dosung Sohn, Andrea B. Kohn, Emilie Neveu, Frederique Varoqueaux, Dirk Fasshauer

## Abstract

Nitric oxide (NO) is a free radical gaseous messenger with a broad distribution across the animal kingdom. However, the early evolution of nitric oxide-mediated signaling in animals is unclear due to limited information about prebilaterian metazoans such as placozoans. Here, we analyzed NO synthases (NOS) in four different species of placozoans (haplotypes H1, H2, H4, H13). In contrast to all other invertebrates studied, *Hoilungia* and *Trichoplax* have three distinct NOS genes, including PDZ domain-containing NOS. To characterize NOS activity in *Trichoplax adhaerens,* we used capillary electrophoresis for microchemical assays of NO-related metabolites. Specifically, we quantified nitrites (products of NO oxidation) and L-citrulline (co-product of NO synthesis from L-arginine), which were affected by NOS inhibitors confirming the presence of functional NOS. Next, using fluorescent single-molecule *in situ* hybridization, we showed that distinct NOSs are expressed in different subpopulations of cells, with a noticeable distribution close to the edge regions of *Trichoplax.* These data suggest the compartmentalized release of this messenger and a greater diversity of cell types in placozoans than anticipated. We also revealed a dramatic diversification of NO receptor machinery, including identification of both canonical and novel NIT-domain containing soluble guanylate cyclases as putative NO/nitrite/nitrate sensors. Thus, although *Trichoplax* is considered to be one of the morphologically simplest free-living animals, the complexity of NO-cGMP-mediated signaling is greater to those in vertebrates. This situation illuminates multiple lineage-specific diversifications of NOSs and NO/nitrite/nitrate sensors from the common ancestor of Metazoa.

**Short Abstract:** Nitric oxide (NO) is a ubiquitous gaseous messenger, but we know little about its early evolution. Here, we analyzed NO synthases (NOS) in four different species of placozoans – one of the early-branching animal lineages. In contrast to other invertebrates studied, *Trichoplax* and *Hoilungia* have three distinct NOS genes, including PDZ domain-containing NOS. Using ultra-sensitive capillary electrophoresis assays, we quantified nitrites (products of NO oxidation) and L-citrulline (co-product of NO synthesis from L-arginine), which were affected by NOS inhibitors confirming the presence of functional enzymes in *Trichoplax*. Using fluorescent single-molecule *in situ* hybridization, we showed that distinct NOSs are expressed in different subpopulations of cells, with a noticeable distribution close to the edge regions of *Trichoplax.* These data suggest both the compartmentalized release of NO and a greater diversity of cell types in placozoans than anticipated. NO receptor machinery includes both canonical and novel NIT-domain containing soluble guanylate cyclases as putative NO/nitrite/nitrate sensors. Thus, although *Trichoplax* and *Hoilungia* exemplify the morphologically simplest free-living animals, the complexity of NO-cGMP-mediated signaling in Placozoa is greater to those in vertebrates. This situation illuminates multiple lineage-specific diversifications of NOSs and NO/nitrite/nitrate sensors from the common ancestor of Metazoa.

## Introduction

Nitric oxide (NO) is a versatile gaseous transmitter widely distributed among prokaryotes and eukaryotes^1–4^ Multiple functions of this messenger are direct reflections of the free-radical nature of NO and, subsequently, its complex free radical chemistry^5^. Dissolved NO passes readily across membranes and diffuses into neighboring cells interacting with many biological molecules including DNA, lipids, proteins^5^ with several specialized receptors such as guanylate cyclases^6–8^. Thus, NO can act as a volume transmitter locally, and it is easily converted into nitrite and nitrate radical by oxygen and water. In cells, NO is catalyzed by the enzyme NO synthase (NOS) through a series of complex redox reactions by the deamination of the amino acid L-arginine to L-citrulline. The reaction requires the presence of oxygen as a precursor and NADPH^5^. The large enzyme operates as a dimer and consists of two enzymatic portions, an oxygenase domain that binds heme and the redox factor tetrahydrobiopterin (H4B) and a reductase domain that is related to NADPH-dependent microsomal cytochrome P450^9^

The role and mechanism of NO signaling are well studied in mammals. However, little is known about the early evolution of NO signaling in animals, mostly due to limited comparative data from basally branching metazoans, including Cnidaria, Porifera, Ctenophora, and Placozoa.

Among other things, NO is involved in feeding, chemosensory processing, and locomotion of such cnidarians as *Hydra* and *Aglantha*^10–13^, where NO-dependent communications were likely mediated by just one type of NO synthase (NOS)^1^. In the sponge *Amphimedon,* only one NOS gene has been identified^14^ NO-cGMP signaling has been implemented in the regulation of larval settlement^15^ and rhythmic body contractions ^16^. In the ctenophore, *Mnemiopsis leidyi,* again, only one NOS gene has been recognized so far^17^, but the functional role of NO has not been studied. Interestingly, in another ctenophore species, *Pleurobrachia bachei,* NOS appears to have been lost^18^.

Nothing is known about the presence and the distribution of NO signaling in Placozoa – an important but little-studied lineage of cryptic marine animals. The current consensus stands that Placozoa is the sister group to the clade Cnidaria+Bilarteria^18–20^, although some authors consider Placozoa as highly derived and secondarily simplified cnidarians^21^. Regardless of the proposed phylogenies, Placozoa represents a crucial taxon to understand the origin and evolution of animal traits and the nervous system in particular^22^

Placozoans, such as *Trichoplax* and their kin^23^, are the simplest known free-living animals with only six morphologically recognized cell types organized in three layers^24^ Nevertheless, *Trichoplax* has quite complex behaviors^25–28^, including social-like patterns^29^ Here, we biochemically showed that *Trichoplax* exhibits functional NOS activity, and, in contrast to other pre-bilaterian animals, placozoa independently evolved three distinct NOSs (as vertebrates) with a profound diversification of NO-cGMP signaling components, and likely the capabilities of nitrite/nitrate sensing by distinct NIT domain-containing guanylyl cyclases, which represents a remarkable example of the evolution of gaseous transmission in the animal kingdom.

## Materials and Methods

### Animals and culturing

*Trichoplax adhaerens* (H1 haplotype) and *Hoilungia hongkongensis* (H13 haplotype), 0.3-2 mm in diameter, were maintained in the laboratory culture as described elsewhere, and animals were fed on rice grains and algae^24,30^.

Direct microchemical assays of NOS metabolites such as NO_2_^-^, L-arginine, L-citrulline were performed using high-resolution capillary electrophoresis (CE) with both conductivity and laser-induced fluorescence (LIF) detectors. The principles and details of major protocols for NOS activity assay were reported^31–33^ with some minor modifications. We made minor adjustments to these protocols, which we briefly summarize below.

### Nitrite/Nitrate Microanalysis using CE with Contactless Conductivity

CE, coupled with a TraceDec contactless conductivity detector (Strasshof, Austria) was used for the assay of nitrite and nitrate. To reduce Cl^-^ in a sample, we used OnGuard II Ag (DIONEX Corp., Sunnyvale, CA). We used custom-built cartridges for small volume (20 μL) sample clean-up by a solid-phase extraction technique as reported^34^. In brief, 4~5 mg of the resin was backloaded in a 10 μL filter-pipette tip, and the microcartridge was washed with 1 mL of ultrapure water using a 3 mL disposable syringe. The pre-washed cartridge was put into a 200 μL pipette tip to avoid surface contamination during further centrifugation. Extra water remaining in the cartridge was removed by centrifugation at 1000 rpm for 30 seconds. Then, the assembly was inserted into a 0.5 mL PCR tube, and a final diluted sample was loaded into the preconditioned cartridge followed by centrifugation at 1000 rpm for 30 seconds, causing the sample to pass through the silver resin. To quantitate any potential sample loss, the custom-made chloride cartridge was tested for sample recovery of both nitrite and nitrate.

All experiments were conducted using a 75 cm length of 50 μm I.D. × 360 μm O.D. fused silica capillary (Polymicro Technologies, AZ) with an insulated outlet conductivity cell. Arginine/borate electrolyte was used for a separation buffer with tetradecyltrimethylammonium hydroxide (TTAOH) added as an EOF modifier. The modifier was prepared from tetradecyltrimethylammonium bromide (TTABr) by an OnGuard-II A cartridge (DIONEX Corp., CA) treated with 1 M NaOH. For separation steps, the capillary inner-wall was successively washed with 1M NaOH, ultrapure water, and the separation buffer (25 mM Arg, 81 mM Boric acid, and 0.5mM TTAOH, pH 9.0) by applying pressure (1900 mbar) to the inlet vial. Since nitrite and nitrate concentrations were very low in diluted samples, capillary isotachophoresis (CITP), a sample stacking method, was employed. The leading solution was introduced into the capillary by pressure injection (25 mbar for 12s), and then a neuronal sample was loaded using electrokinetic injection (−5kV for 12s). The separation was performed under a stable – 15kV voltage at 20°C.

### Amino Acids Microanalysis using CE with laser-induced fluorescence detection

The CE, coupled with the ZETALIF detector (Picometrics, France), was used for the assay of amino acids. In this work, a helium-cadmium laser (325nm) from Melles Griot, Inc. (Omnichrome^®^ Series56, Carlsbad, CA) was used as the excitation source. Before the photomultiplier tube (PMT), the fluorescence was both wavelengths filtered and spatially filtered using a machined 3-mm pinhole. All instrumentation, counting, and high-voltage CE power supply were controlled using DAx 7.3 software.

All solutions were prepared with ultrapure Milli-Q water (Milli-Q filtration system, Millipore, Bedford, MA) to minimize the presence of impurities. Borate buffer (30 mM, pH 9.5) was used for sample preparation. All solutions were filtered using 0. 2 μm filters to remove particulates. The buffers were degassed by ultrasonication for 10 min to minimize the chance of bubble formation. A 75 mM OPA/β-mercaptoethanol (β-ME) stock solution was prepared by dissolving 10 mg of OPA in 100 μL of methanol and mixing with 1 mL of 30 mM borate and 10 μL of β-ME. Stock solutions (10 mM) of amino acids were prepared by dissolving each compound in the borate buffer. OPA and β-ME were stored in a refrigerator, and fresh solutions were prepared weekly.

All experiments were conducted using a 75 cm length of 50 μm I.D. × 360 μm O.D. fused silica capillary (Polymicro Technologies, AZ). A 30 mM borate/30 mM sodium dodecyl sulfate (SDS) electrolyte (adjusted to pH 10.0 with NaOH) was used as the separation buffer for amino acid analysis. The pre-column derivatization method was used. A 1 μL of o-Phthalaldehyde (OPA) was incubated in a 0.5 mL PCR tube. The total volume of a sample, OPA, and internal standard inside the tube was 20 μL. For separation steps, the capillary inner-wall was successively washed with 1M NaOH, Milli Q water, and the separation buffer by applying pressure (1900 mbar) to the inlet vial. Then the sample was loaded using electrokinetic injection (8 kV for 12s). The separation was performed under a stable 20 kV voltage at 20°C.

In all CE tests, once an electropherogram was acquired, peaks were assigned based on the electrophoretic mobility of each analyte, and the assignments were confirmed by spiking corresponding standards into the sample. Five-point calibration curves (peak area vs. concentration) of analytes were constructed for quantification using standard solutions. All chemicals for buffers were obtained from Sigma-Aldrich, and standard amino acids were purchased from Fluka. Ultrapure Milli-Q was used for all solutions and sample preparations.

### NOS inhibitors’ tests

To establish that NOS enzymatic activity is responsible for producing the Arg/Cit ratio and nitrite measured in *Trichoplax*, the whole animal was incubated in one of NOS inhibitors (e.g., *N*^G^-nitro-l-arginine methyl ester (L-NAME); besides, another NOS inhibitor, L-N^6^-(1-iminoethyl)-lysine (L-NIL), showed very effective inhibition as in molluscan preparations^35^.

After the animals were isolated from the culture medium, they were placed in a 0.5 mL PCR tube and incubated with certain concentrations of NOS inhibitors for 30 minutes at room temperature, followed by washing with artificial seawater. Then, all the water was removed, and 1 μL of Milli Q water was dropped onto the animal, and the tube was stored at −80° C until use.

Specifically, we also performed a series of control tests to see if there were any small molecules that might interfere with peak identifications. Water, L-NAME, and L-NIL controls were first tested, and no nitrite was observed. However, chloride and nitrate ions were always present, because all NOS inhibitors contain chloride, and nitrate is a common impurity in most of the commercially used chemicals. Fresh single individuals of *Trichoplax* by itself, and *Trichoplax* incubated with NOS inhibitors were then analyzed. An effective NOS inhibitor should cause the nitrite level to be lower than in the animal compared to control tests.

### Comparative bioinformatic analyses

We used the data from the sequenced genomes of two sequenced placozoan species^36,37^, and our additional sequencing data are presented in the supplement 1.

Protein sequences were aligned in MUSCLE^38^. Phylogenetic trees were inferred using Maximum Likelihood algorithm implemented in IQTREE web server http://iqtree.cibiv.univie.ac.at/,^39^. Tree robustness was tested with 2000 replicates of ultrafast bootstrap^40,41^.

To test for positive and negative selection, the following algorithms were used: codon-based Z-test and Fischer’s exact test implemented in MEGA X^42–44^, and ABSREL, BUSTED, FUBAR and MEME in HyPhy package^45–49^. Evolutionary distances were calculated in MEGA X under the Poisson method and gamma-distributed rates across sites.

**Fixative-resistant NADPH-diaphorase activity** has been widely used as a histochemical reporter of NOS in both vertebrates and invertebrates^33,50–53^. Thus, we used this approach to screen for putative NOS activity in *Trichoplax* and *Hoilungia*. All methodological details of the protocol have been described earlier^54–57^, and we used 45 and 90 min fixation in 4% freshly made paraformaldehyde solution made using the filtered seawater.

***In situ* hybridization** was performed using the RNAscope multiplex fluorescent Reagent kit v2 assay (Advanced Cell Diagnostics, Inc, Bio-Techne, USA) as specified by the company protocol (https://acdbio.com/rnascope%C2%AE-multiplex-fluorescent-v2-assay). In brief, we transferred 10-15 animals to the glass slides with cavities with 2 mL fresh 0.2 μm filtered seawater, washed three times, and removed the seawater under a microscope. Next, we fixed animals using 4% paraformaldehyde in seawater for 30 min at room temperature, performed dehydration and rehydration steps with increased and decreased concentrations of ethanol (30%, 50%, 70%, 100% on PBS) at room temperature. We pretreated animals in Protease III (Sigma) for 10 min at room temperature. The rest of the protocol is reported elsewhere (Advanced Cell Diagnostics, ACD #323110 at web site https://acdbio.com/rnascope%C2%AE-multiplex-fluorescent-v2-assay). The key point in the procedure is to use tyramide signal amplification steps to detect low – abundant genes as NOSs.

For all imaging, we used fluorescent microscope Nikon Ti2 (Nikon, Japan) with a spinning disk (Crest Optics X-Light V2).

## Results and Discussion

### Comparative analysis of NOSs

Fig. 1 shows the genealogical relationships among different animal NOSs, where representatives of all basal metazoan lineages form distinct branches for their respected NOSs with evidence for relatively recent duplication events consistent to an early origin and diversification of NOSs in other eukaryotic groups including Amebozoa and Fungi as sister lineages to Metazoa. We did not find the evidence for NOS in choanoflagelates sequenced so far, including *Monosiga* with the sequenced genome. Choanozoa, the phylogenetically closest taxon to Metazoa^58^, might have lost NOS from its eukaryotic ancestor. Interestingly, ctenophores, known as the sister lineage to the rest of metazoans^19,20,59^, have only one highly derived NOS as represented by two *Mnemiopsis* species and *Cestum* in the tree (Fig. 1).

**Figure 1.**
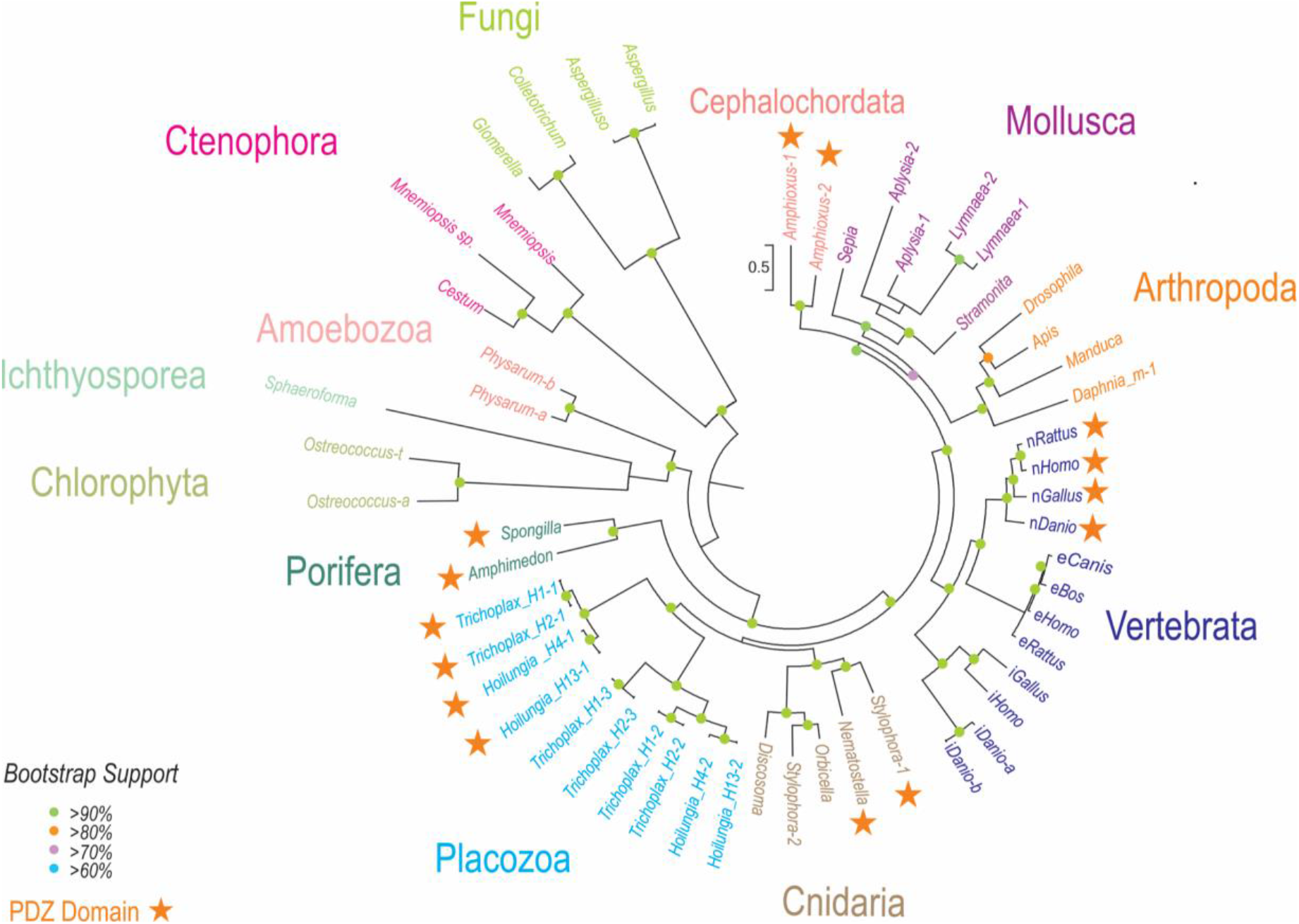
The diversity and evolutionary relationships of nitric oxide synthases in animals. The representative lineages of deuterostomes, protostomes, and basal metazoans are highlighted with unicellular eukaryotes and algae as outgroups. Names of the species are indicated in each case with the existing classification of NOSs (see text for details). The references for each particular gene with relevant gene accession numbers are summarized in the method section and supplementary Table 1.

All studied invertebrates have only one or two NOS genes, which do not directly correspond to the well-established vertebrate subfamilies of the enzymes^1^. In contrast, we identified three distinct NOS genes in the *Trichoplax* genome (haplotype H1), and three other placozoan species or haplotypes H2, H4, H13, and one of the NOSs contains the PDZ domain similar to the mammalian neuronal NOS.

The presence of PDZ domain-containing NOSs is a distinct feature of all four placozoan species sequenced so far (NOS1 in H1, H2, H4, and H13). H1 and H2 represent the classical *Trichoplax* genus^23^, while H4 and H13 belong to the newly described genus *Hoilungia*^36,60^. The clustering of NOSs in placozoans reflects their phylogenetic relationships stressing that H4 and H13 (*Hoilungia*) vs. H1 and H2 *(Trichoplax)* belong to different lineages.

The PDZ domain and N-terminal motifs are required for the anchoring of NOS to plasmatic or intracellular membranes, subcellular localization, and integration to many signaling components like in the mammalian neuronal nNOS^9,61–64^ nNOS is different from the two other mammalian isoforms as its N-terminal PDZ domain can heterodimerize with the PDZ domains of PSD95 or syntrophin^65^ and others^9^ Thus, we might suggest similar molecular functions in *Trichoplax* and *Hoilungia*.

The rate of evolution of the PDZ domain-containing NOSs is comparable to other NOSs for all placozoan species. The branching patterns of NOS trees (Fig. 1) reveals that three NOSs in Placozoa are the results of two independent duplication events from the common placozoan ancestor. The first splitting separated NOS1, and the second, more recent split led to NOS2 and NOS3.

Of note, we also identified two NOSs in the stony coral *Stylophora*, which has one NOSs with the PDZ domain (Fig. 1), and a PDZ domain was detected upstream of the *Nematostella* NOS gene. Also, two sponges *(Amphimedon* and *Spongilla)* possess PDZ-containing NOS. As the PDZ domains of NOSs appear to be homologous, it should be investigated whether the PDZ domain-containing form represents the ancestral form in animals. In contrast, ctenophores and many other animal lineages (Fig. 1) do not have PDZ-containing NOS genes.

NOS is a complex enzyme requiring several co-factors for its activation, and Ca^2^+-dependence of different NOSs in mammals is determined by the presence of the autoinhibitory inserts and calmodulin-binding sites^66–73^. Fig. 2 shows the presence/absence of such motifs and the auto-inhibitory loops across basal metazoan lineages. The canonical human Ca^2+^-independent iNOS lacks such a loop; it is bound to CaM in a Ca^2^+-independent manner. Mammalian iNOS activation is often induced by lipopolysaccharides as a part of innate immunity responses on bacterial infection^4^. Ctenophore, the demosponge *Amphimedon*, *Nematostella,* and three coral NOSs also lack the auto-inhibitory loop (Fig. 2) and could be Ca^2+^-independent and, apparently, inducible (e.g., by bacteria or during development and differentiation). However, all three *Trichoplax* and *Hoilungia* NOSs contains an intermediate size insert in this position: these NOSs might be dormant or, at least, partially inducible. Thus, the direct detection of endogenous enzymatic activity is needed to validate NOS expression, which we performed using direct microchemical assays.

**Figure 2.**
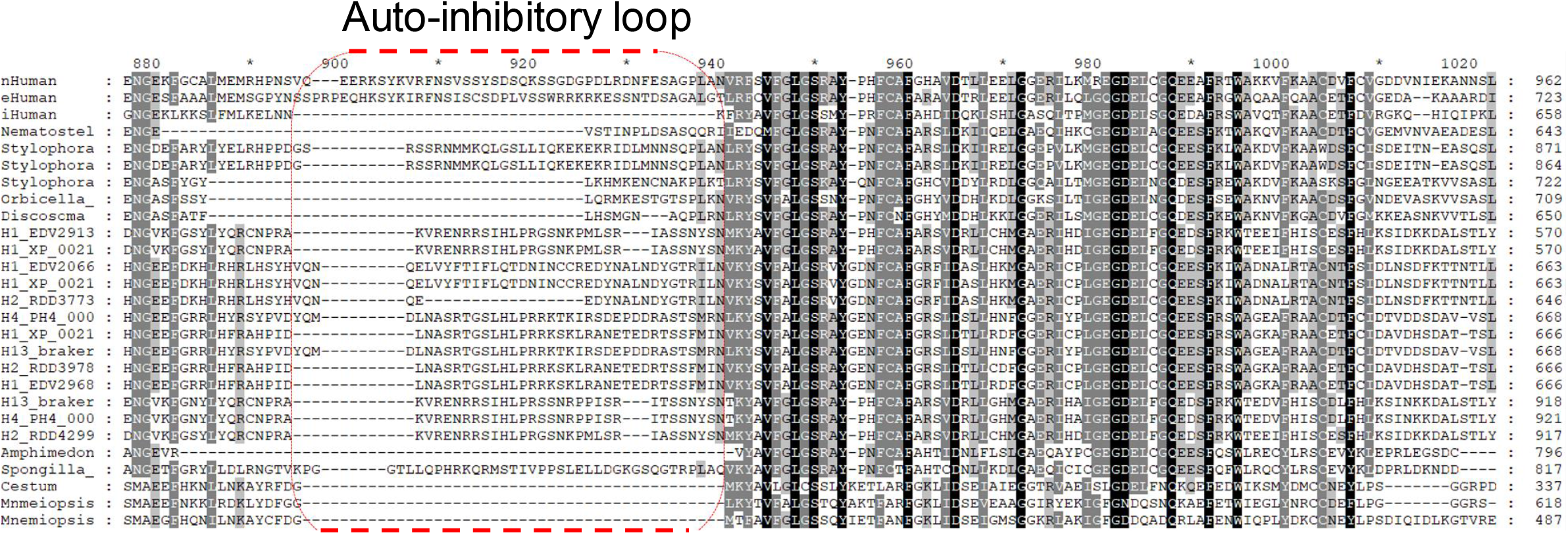
Auto-inhibitory N-terminal inserts in NOSs: Insights into controlling Ca^2+^-dependence and the conservation of calmodulin-binding sites in placozoans. Alignments of selected NOSs was performed using the sequence information from species outlined in Fig. 1.

### Detection of endogenous NOS activity in *Trichoplax*

Because some NOSs can be inducible or pseudogenes, the molecular/sequence information itself is not sufficient for the demonstration of NOS activity. Thus, we implement two complementary approaches to confirm the presence of functional NOSs in placozoans.

#### Arginine/citrulline assays

NO is known to be produced enzymatically from molecular oxygen and L-arginine with L-citrulline as the co-product^5^. Using a highly sensitive capillary electrophoresis (CE) microchemical assay with attomole detection limits, we demonstrated that *Trichoplax* produced L-citrulline, and its production is also eliminated by NOS inhibitors (Fig. 3). It was expected from experiments on vertebrates and mollusks^32,33,35^ that the arginine-to-citrulline ratio would increase after *Trichoplax* was incubated in either L-NAME/D-NAME or L-NIL. the arginine-to-citrulline ratio increased by two-fold in the case of L-NIL (Fig. 3). However, there was only a small increase with L-NAME, indicating L-NIL effectively inhibited the NOS enzyme as in mollusks^32,33,35^, but L-NAME did not. The reason for this difference might reflect differences in either L-arginine uptake, which might be blocked by arginine analogs or distinct enzymatic regulation of NOS in placozoans, or nonenzymatic interference of these inhibitors with NO production^74^.

**Figure 3.**
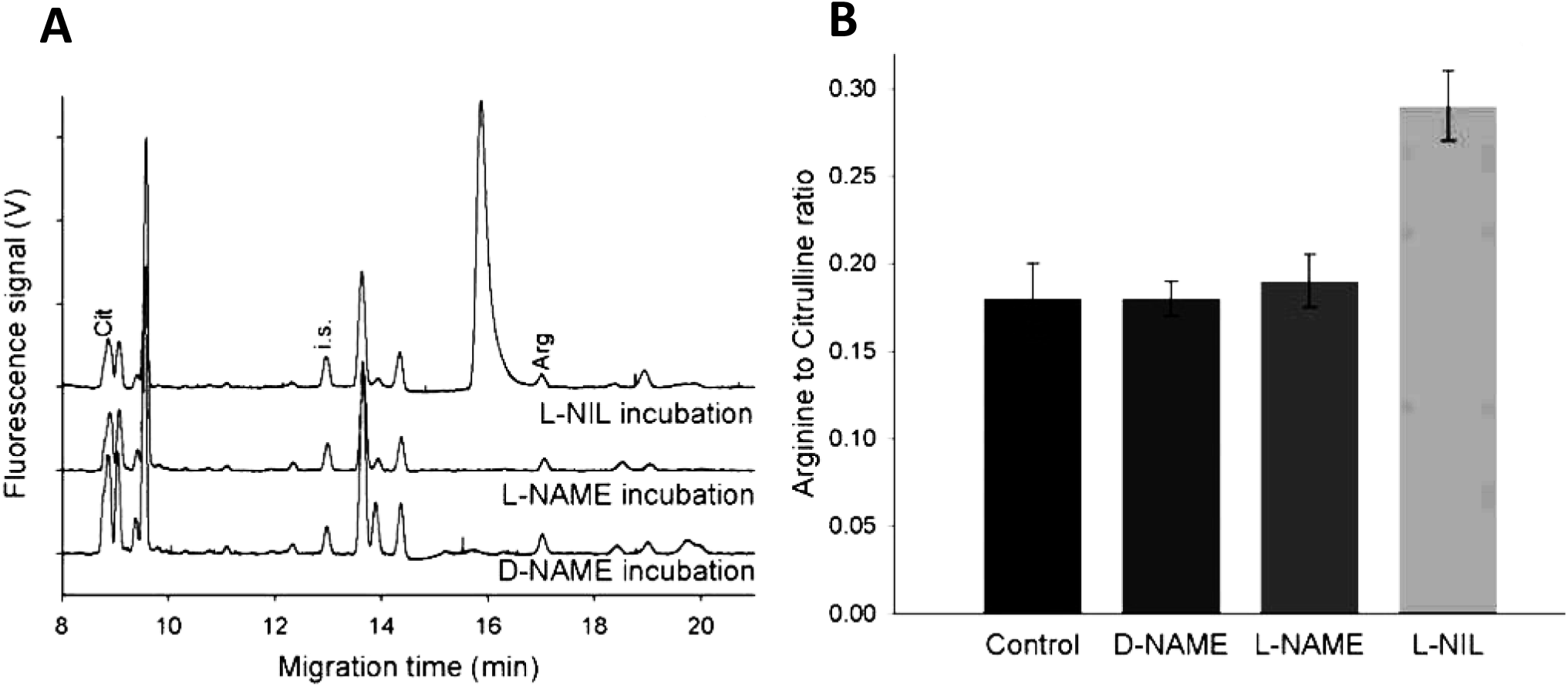
Detection of NOS amino acid-derived metabolites by capillary electrophoresis and their sensitivity to NOS inhibitors. **A**. Electropherograms of individual animal samples and L-Arginine to L-Citrulline ratios of *Trichoplax adhaerens* following treatment with NOS inhibitors. Arginine and citrulline peaks were identified with spike standards and shown as Arg and Cit, respectively. i.s. is internal standards (see methods for details). Samples were loaded using electrokinetic injection (8 kV for 12s) and then analyzed under a stable 20kV voltage at 20°C in 50 μm I.D. and 360 μm O.D. capillary with 30 mM borate/30 mM SDS, pH 10.0. (A) Electropherograms of *Trichoplax* incubated with *N*^G^-nitro-l-arginine methyl ester or L-NAME (500 μM), D-NAME (500 μM), and L-N^6^-(1-iminoethyl)-lysine, L-NIL (1 mM), for 30 min at room temperature. **B.** Arginine-to-Citrulline ratio of *Trichoplax* after treatment with putative NOS inhibitors; only L-NIL induced statistically significant increase of Arg/Cit ration suggesting the suppression of L-citrulline production (n=5, p < 0.05, see results for details).

It was interesting that all NOS-related metabolites were detected in *Trichoplax* at relatively high concentrations, 0.35 mM for arginine and 0.5 mM for citrulline. Combined, these CE/microchemical data indicate that placozoans have a substantial level of endogenous NOS activity.

#### Nitrite assays

Due to rapid NO oxidation in biological tissues^5^, NO_2_^-^ is considered as the most reliable reporter of functional NOS. In contrast, more stable (and less dynamic) terminal oxidation products of NO – nitrates (NO_3_^-^) cannot be used for these purposes since they can also be accumulated from various food sources. Thus, by employing CE with the conductivity detection, we provided the additional direct evidence for endogenous NOS activity using nitrite (NO_2_^-^) assay^31,33^.

NO oxidation metabolites were monitored, and concentrations were derived from *in vitro* calibration curves prepared from standard solutions of nitrate and nitrite at various concentrations (10 nM – 500 μM). With the regression equations, the limit of detection (LOD) of nitrate was determined to be 13.3 nM for nitrite and 32.4 nM for nitrate. These LODs were sufficient to quantify nitrite and nitrate in *Trichoplax*.

Surprisingly, we found than *Trichoplax* contains millimolar concentrations of NO_2_^-^ and NO_3_^-^, which were eliminated, within 30 min, by NOS inhibitors such as L-NAME and L-NIO (Fig. 4). In control *Trichoplax*, about 1.5 mM nitrite was detected, but after incubated with the NOS inhibitors, no nitrite was observed, suggesting the suppression of endogenous NOS activity (Fig. 4).

**Figure 4.**
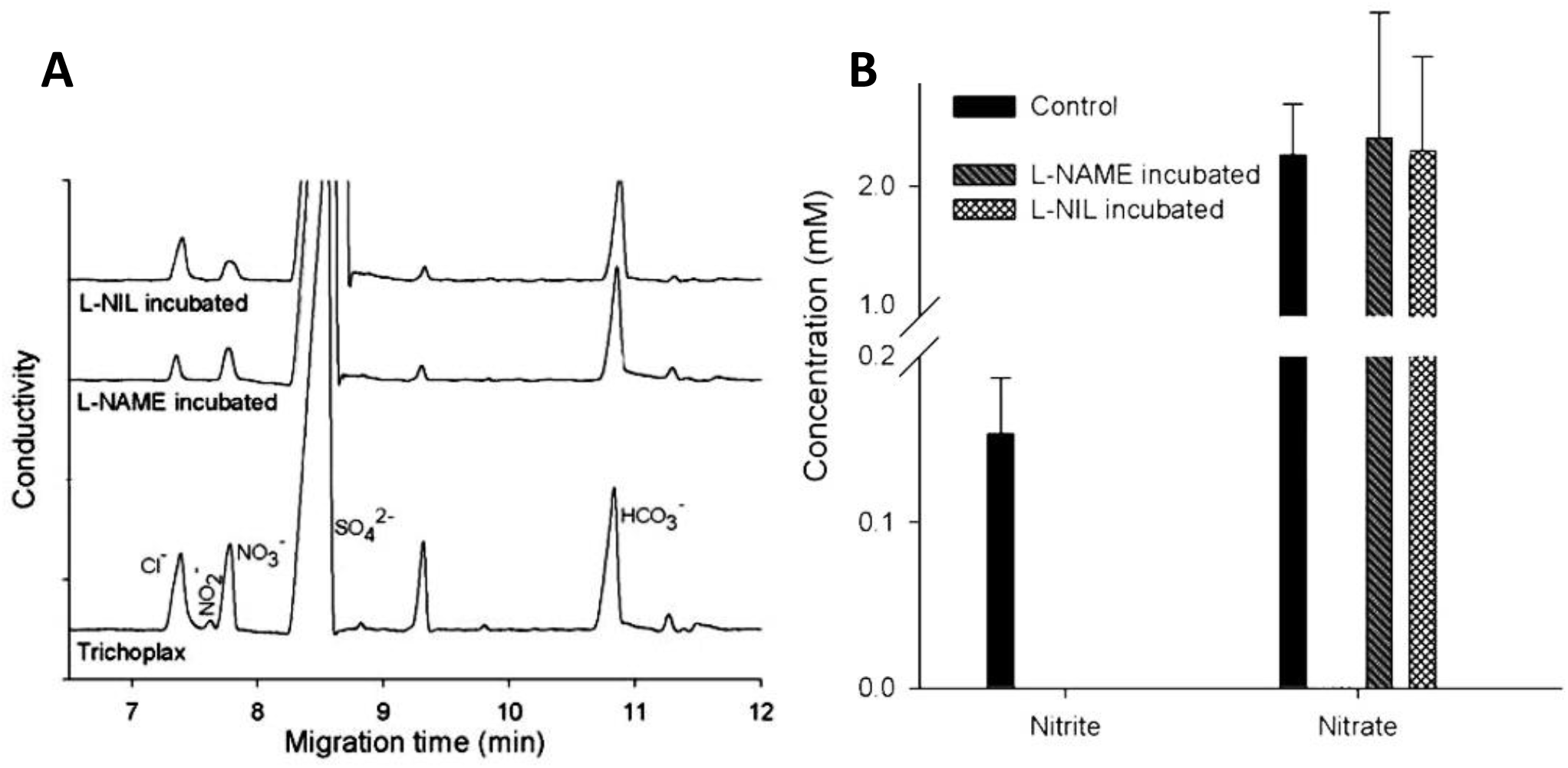
Detection of products of NO oxidation (NO_2_^-^ and NO_3_^-^) by capillary electrophoresis and their sensitivity to NOS inhibitors. Nitrites, products of NO oxidation, were detected in all control samples and eliminated following NOS inhibitor incubation (see text for details). The separation was conducted in a 75cm length of 50 μm I.D. and 360 μm O.D. capillary with arginine/borate buffer, pH 9.0. All samples were loaded using electrokinetic injection (−1 kV for 12s), and then analyzed under a stable −15 kV voltage at 20°C. **A**. Electropherograms of *Trichoplax* only, and *Trichoplax* incubated for 30 mins with *N*^G^-nitro-l-arginine methyl ester or L-NAME (500 μM), and L-N^6^-(1-iminoethyl)-lysine or L-NIL (1 mM). **B**. Nitrite and nitrate concentration profiling after 30 mins of NOS inhibition (n=5, p < 0.05).

### The expression and distribution of NOS in *Trichoplax*

Fixative-resistant NADPH-diaphorase (NADPH-d) histochemistry has been reported as a marker of functional NOS in both vertebrates and invertebrates^33,50–53^. Here, we employed this assay for the initial screening of the NOS expression in *Trichoplax adhaerens* (H1) and its related species *Hoilungia hongkongensis*^36^. The NADPH-d histochemical activities in both placozoans were significantly weaker compared to the majority of other species studied using the same protocol^13,54,55,57,75–77^ We noted that the intensity of NADPH-d labeling was similar to those described in the pelagic pteropod mollusk *Clione limacina,* where NO controlled swimming^56^.

We revealed very similar NADPH-d labeling patterns in both *Trichoplax* and *Hoilungia* (Fig. 5A, B). There were several large (>10 μm) structures; some of them correspond to the so-called “shinny spheres”^78^ and numerous small (4-6 μm) NADPH-d reactive cells were broadly distributed over different parts of the animal including the dorsal epithelial layer. We estimate that about 2% of placozoan cells might be NADPH-d reactive. These cells might be candidates for NOS-containing (NO-releasing) cells. However, NADPH-d histochemistry cannot distinguish different NOS isoforms.

**Figure 5.**
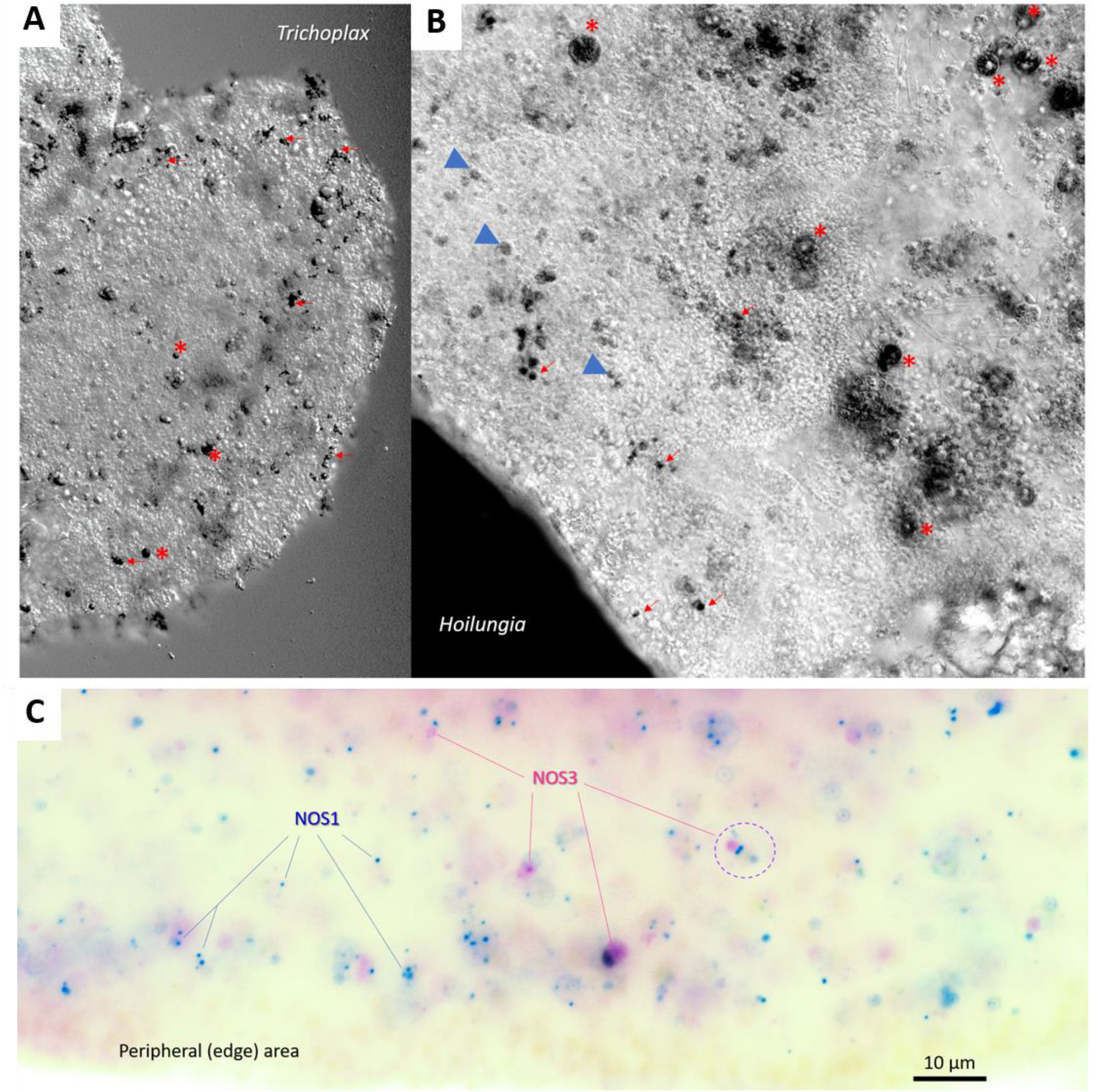
NOS expression in Placozoans. (A, B) NADPH-diaphorase histochemistry and the distribution of putative nitrergic cells in two species of Placozoa: *Trichoplax adhaerens* (**A**) and *Hoilungia hongkongensis* (**B**). NADPH-d reactive cells (black) are broadly distributed across the animal. In both species, relatively large cells (asterisks) correspond to so-called “shiny spheres,” whereas the arrows indicate an example of NADPH-d reactive cells with some tendencies of their distribution close to the edge of animals. (**C**) Expression of two NOSs in *Trichoplax* using single molecules fluorescent *in situ* hybridization (FISH). Blue dots – PDZ-containing NOS1 and purple dots – NOS3. A dotted circle indicates an example of a cell where both NOS are co-localized. Note, NOS-expressed cells do not occur at the very edge of the animal. Scale: 10 μm.

### Single-molecule in fluorescent *in situ* hybridization (FISH)

Next, we used sequences for both NOS1 and NOS3 to characterize their expression and distribution in *Trichoplax adhaerens* by single-molecule FISH as the most sensitive assay for this purpose. In both cases, we observed the cell-specific distribution of distinct NOS isoforms (Fig. 5C). Most of the NOS-containing cells were broadly distributed (similar to NADPH-d reactivity, but ‘shiny spheres” were not labeled by *in situ* hybridization probes). It appears that PDZ containing NOS1 expressed in more cells than NOS3, and only partial co-localization of the two NOSs was observed (Fig. 5C). We also noted that the NOSs are not located to the most peripheral cell layer but found in cells close to the edge. Due to a relatively high level of endogenous fluorescence in the central part of the animal, the precise cell identity of NOS-positive cells was difficult to determine. However, we noticed that both NOS could be co-localized in a very small subset of cells close to the edge of these disk-like animals.

### NO targets and diversification of cGMP signaling in Placozoa

NO can act via cyclic guanosine monophosphate or cGMP as second messenger. In this signaling pathway, NO binds to the heme group of soluble guanylate cyclases (sGCs), member of the adenylyl cyclase superfamily^79,80^, with a characteristic catalytic CYC domain, leading to the increase of cGMP synthesis^6–8,81–85^; by binding to ATP, sGC can also couple NO signaling to cellular metabolism^86^.

Surprisingly, the *Trichoplax* and *Hoilungia* genomes encode seven sGCs (Fig. 6A), whereas only three orthologs were identified in humans. All these enzymes have the canonical heme NO binding domain and associated domain organization, and the predicted sGCs from placozoans from clusters appropriately with the α and β sGCs of humans.

**Figure 6.**
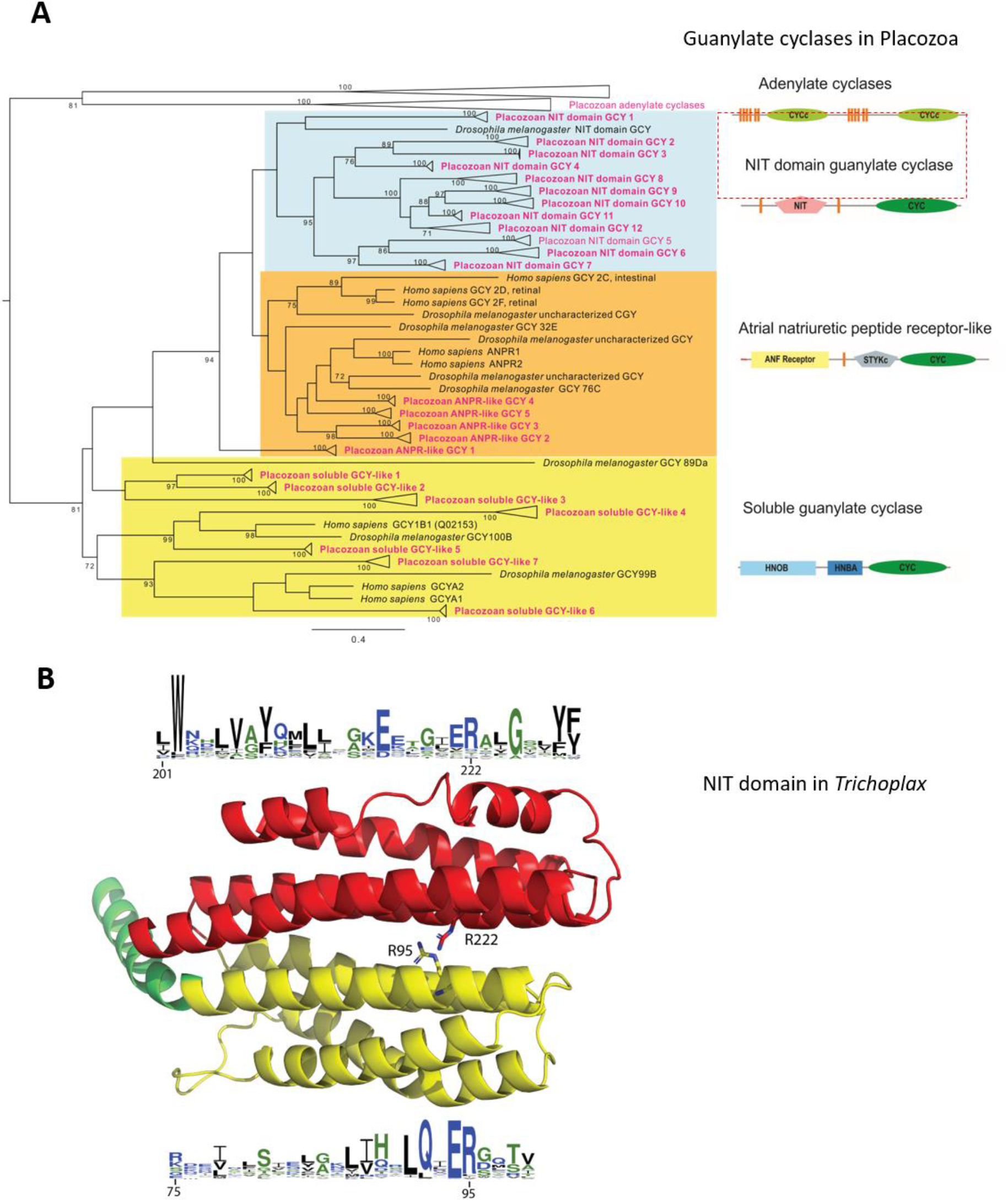
The diversity and lineage-specific expansion of sGC and related NO receptors in placozoans. **A.** Maximum likelihood phylogenetic tree of placozoan soluble guanylyl cyclases (sGC) and two groups of related enzymes: Atrial Natriuretic Peptide-like receptors (ANPRs), some of which contain unusual NIT domains – putative nitrite/nitrate sensing receptors (see text), and adenylate cyclases as outgroups. 119 protein sequences (Supplementary Table 1) were trimmed down to cyclase domains and produced an alignment 325 aa long. Alignment was analyzed in IQTREE^92^ using LG+I+G4 evolution model chosen automatically with Bayesian information criterion. Tree robustness was tested with 2000 replicates of ultrafast bootstrap. Orthologous proteins from 4 placozoan species (red text) were analyzed and their branches collapsed in the tree: *Trichoplax adhaerens* (H1), *Trichoplax* sp. (H2), *Hoilungia* sp. (H4) and *Hoilungia hongkongensis* (H13), except for adenylate cyclases that were only from *Trichoplax adhaerens,* and NIT domain GCY3 which were found only in *Hoilungia* genus. Human and *Drosophila* orthologs are shown. The domain organization of three groups of predicted guanylyl cyclases in placozoans is also schematically illustrated. Full (uncollapsed) version of this tree can be found in Supplemental Figure 1S. **B. NIT domains in placozoans**. The putative nitrate- and nitrite-sensing NIT domains of animals are homologous to prokaryotic NIT domains. Phyre2 was used to generate a structural model for the NIT domain of 007393 from *Trichoplax adhaerens.* The Phyre model is mostly based on the structure NIT domain of the NasR transcription antiterminator (pdb ID: 4AKK). The NIT domain consists of two four-helix bundles, shown in yellow and red. At their interface, two conserved arginines are thought to be involved in ligand binding. The sequence conservation of the two helices at the interface is shown by a webLogo representation^93^. The overall height of a stack indicates the sequence conservation at a certain position, whereas the height of symbols within the stack indicates the relative frequency of each amino acid at that position.

We identified in *Trichoplax* and their kin additional membrane-bound NO receptor candidates (Fig. 6A). *Trichoplax* also has five orthologs of atrial natriuretic peptide-like receptors (ANPRs) with CYC/cGMP coupling as in humans. But there is no atrial natriuretic peptide detected in any sequenced placozoan genome. There are also four *Trichoplax* adenylate cyclases, which have two CYC domains (humans have nine adenylate cyclases); these are probably not involved in NO binding and we used them as outgroups.

Unexpectedly, we discovered 12 additional guanylyl cyclases with unique NIT domains^87^, which were only previously known from bacteria as nitrate and nitrite sensors^88,89^. Nitrate/nitrite sensing type domain in placozoans (NIT: PF08376) is flanked between two transmembrane domains and a C-terminal guanylate cyclase catalytic domain (AC/GC: PF00211). The same critical amino acid residues that were observed in the bacterial sequences were also present in the predicted placozoan NIT domains. (Fig. 6B). The tree shows clearly that they belong to the ANPR type/group and probably arose by lateral gene transfer into an existing ANPR type, which is established as guanylate cyclase.

The heme-dependent NO sensor HNOBA (PF07701) is also found associated with some of these predicted proteins.

To the best of our knowledge, these types of NIT containing proteins have not been previously characterized in animals. There is no NIT domain detected in the sequenced genomes of ctenophores and sponges. However, the observed NIT abundance in placozoans suggests potential sensing of nitrites and/or nitrates. This hypothesis is consistent with our present finding of the micromolar concentration of nitrites in *Trichoplax.* Because many placozoan cells (e.g., fiber cells) do contain endosymbiotic bacteria, additional levels of intra- and intercellular NO-dependent communications are also highly likely and can be tested in future studies.

Even more interesting, we found NIT-containing GCs across bilaterians including molluscs, annelids, arthropods, priapulids, echinoderms, hemichordates and basal chordates but vertebrates apparently lost NIT domains (Fig. 2 Supplement). Apparently, molluscs, hemichordates (*Saccoglossus*) and placozoans have one of the largest numbers of predicted NIT domain genes compared to all studied metazoans.

The model cnidarian *Nematostella* has no NIT domain, but there are NIT-containing genes in the genome of related anthozoan species including corals. The supplementary phylogenetic tree shows that all metazoan NIT-GCs cluster together and their NIT domains are more similar to each other than to bacterial NITs.

The exact function of the NIT domain in animals is yet to be elucidated, but the same architectural domain organization of the NIT domain^89,90^ is observed across metazoans (Fig. 6B) inferring a similar function. In bacteria, it has been proposed that the NIT domain regulates cellular functions in response to changes in nitrate and/or nitrite concentrations, both extracellular and intracellular^88,89^. The same possibility of nitrite/nitrate sensing might be widespread across the animal kingdom. Functional studies would be needed to carefully test this hypothesis in the future.

### Highlights

1. The phylogenetic position of Placozoa, as an early branching metazoan lineage, and the simplicity of morphological organization emphasizes the importance of *Trichoplax* as one of the key reference species for understanding the origin and evolution of animals and their signaling mechanisms^91^, including NO-/cGMP-mediated signaling.
2. Our combined genomic, molecular, and microchemical analyses strongly indicate the presence of functional NOSs in *Trichoplax,* which is broadly distributed across different cell populations. In contrast to other prebilaterian animals, placozoans independently evolved three different NOS genes, similar to the situation in vertebrates. This relatively recent diversification of enzymes producing gaseous free radical messenger illustrates the parallel development of complex signaling mechanisms in placozoans and implies a much greater complexity of intercellular communications than it was anticipated before.
3. The molecular targets of NO in *Trichoplax* can be seven soluble guanylyl cyclases (sGCs) and five membrane-bound ANP-like receptors. Besides, we identified at least twelve cyclases with unique NIT domains. Placozoans have the largest number of predicted NIT domain genes compared to all studied metazoans. We hypothesize that in placozoans, as in bacteria, the putative NIT domain is used as nitrate/nitrite-sensing due to the high levels of nitrate/nitrites measured in *Trichoplax*.

In summary, although canonical functional NO-cGMP signaling could be a highly conservative feature across Metazoa, the enormous diversity of molecular components of these and related pathways in placozoans stress the cryptic complexity of these morphologically simplest animals.

## Acknowledgments

This work was supported by the Human Frontiers Science Program (RGP0060/2017), National Science Foundation (1146575, 1557923, 1548121 and 1645219) grants to L.L.M; and the Swiss National Science Foundation (#31003A_182732) to D.F.

We thank the Cellular Imaging Facility (UNIL, Lausanne, Switzerland) for FHL for their excellent support and Friday Harbor Laboratories for their support to collect animals.

## Role of authors

All authors had access to the data in the study and take responsibility for the integrity of the data and the accuracy of the data analysis. ABK, DYR and MAN share authorship equally. Research design: LLM. Acquisition of data: all authors (Molecular data and sequencing analyses ABK, MAN, DYR, EN, DF, LLM; *in situ* hybridization FV, DYR, LLM, DF; Microchemical assays: LLM and DS; NADPH-d: DYR, LLM). Analysis and interpretation of data: all authors. Drafting of the article: LLM. Funding: LLM, DF.

## Supporting Information

**Figure 1S.**
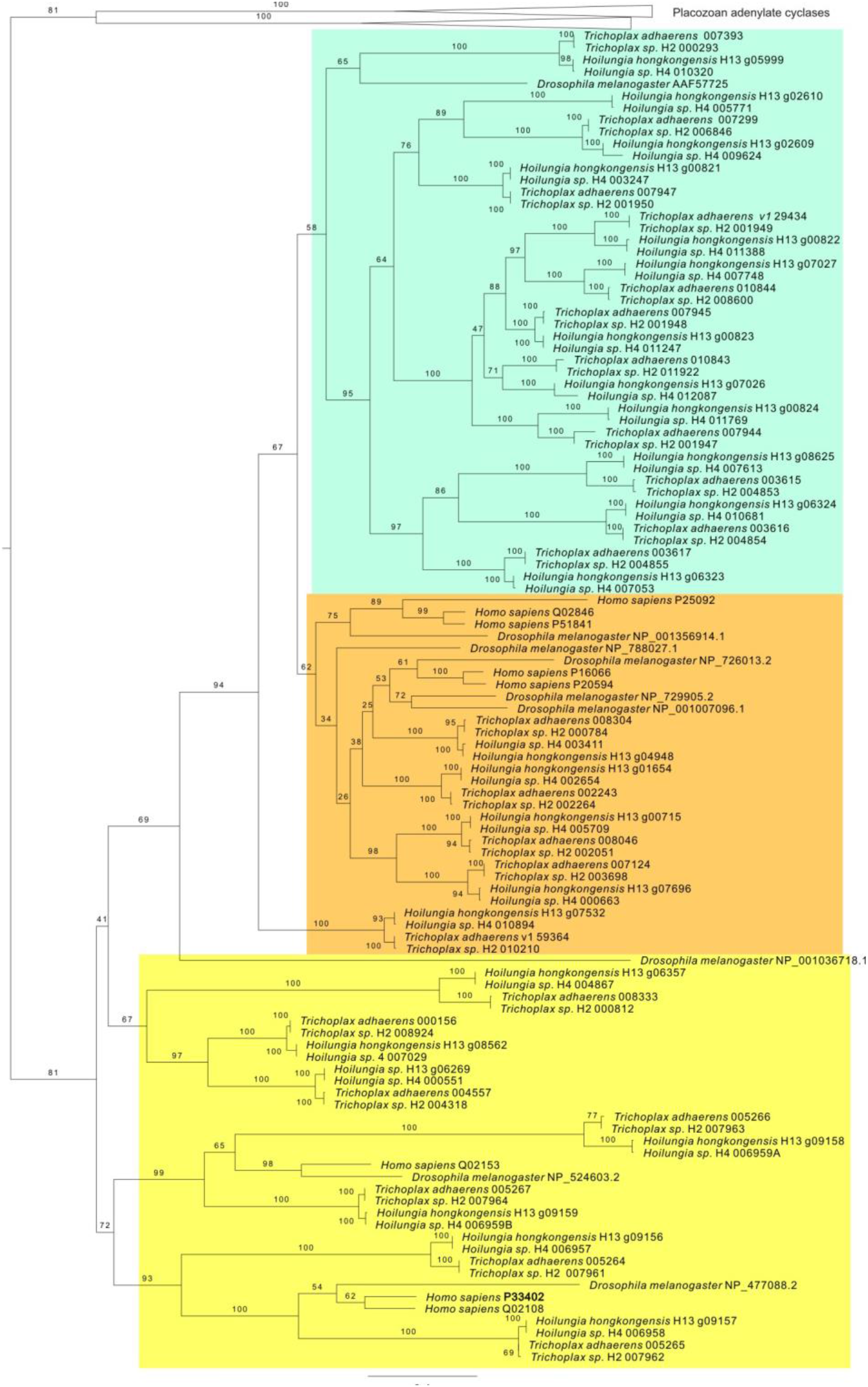
Maximum likelihood phylogenetic tree of placozoan soluble guanylyl cyclases (sGC) and two groups of related enzymes: Atrial Natriuretic Peptide-like receptors (ANPRs), some of which contain unusual NIT domains, and adenylate cyclases. Proteins from all four placozoan species used in this study are represented (see Fig 6A in the main text).

**Figure 2S.**
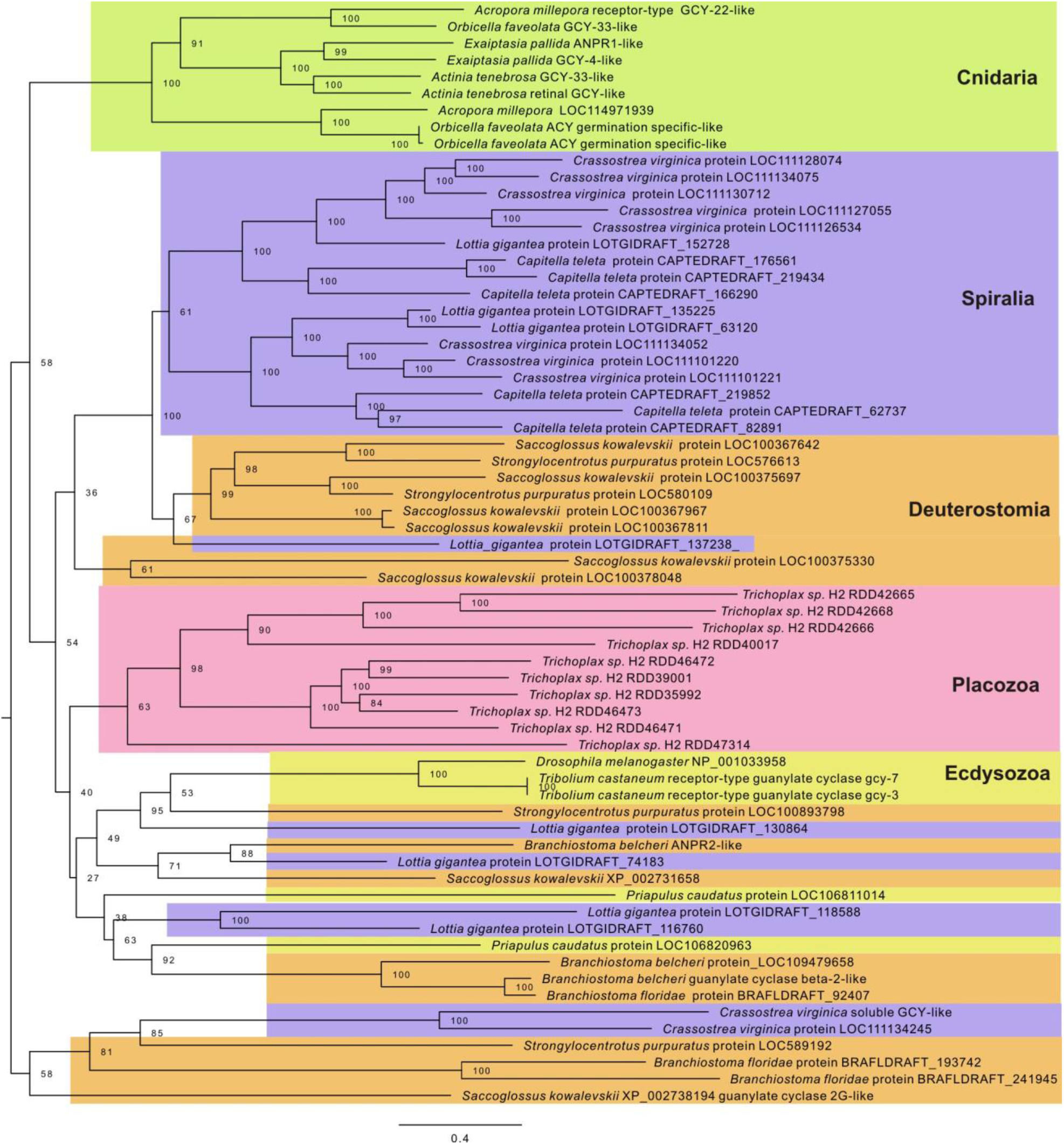
Maximum likelihood phylogenetic tree of NIT domains containing guanylyl cyclases in placozoans *(Trichoplax* sp. H2 only), cnidarians and bilaterians. Guanylate cyclases with NIT domains are found in most animal phyla except sponges, ctenophores, vertebrates and urochordates. Extensive lineage-specific duplications are evident in placozoans, molluscs and hemichordates. 66 protein sequences were trimmed down to NIT+cyclase domains and produced an alignment 689 aa long. Alignment was analyzed in IQTREE^92^ using LG+F+R6 evolution model chosen automatically with Bayesian information criterion. Tree robustness was tested with 2000 replicates of ultrafast bootstrap.

